# Identification and mapping of central pair proteins by proteomic analysis

**DOI:** 10.1101/739383

**Authors:** Daniel Dai, Muneyoshi Ichikawa, Katya Peri, Reid Rebinsky, Khanh Huy Bui

## Abstract

Cilia or flagella of eukaryotes are small micro-hair like structures that are indispensable to single-cell motility and play an important role in mammalian biological processes. Cilia or flagella are composed of nine doublet microtubules surrounding a pair of singlet microtubules called the central pair (CP). Together, this arrangement forms the canonical and highly conserved 9+2 axonemal structure. The CP, which is a unique structure exclusive to motile cilia, is a pair of structurally dimorphic singlet microtubules decorated with numerous associated proteins. Mutations of CP-associated proteins cause several different physical symptoms termed ciliopathies. Thus, it is crucial to understand the architecture of the CP. However, the protein composition of the CP was poorly understood. This was because identification of CP proteins was mostly limited by available *Chlamydomonas* mutants of CP proteins. In this study, we conducted a comprehensive CP proteome analysis using several CP mutants and identified 37 novel CP protein candidates. By using *Chlamydomonas* strains lacking specific CP sub-structures, we also present a more complete model of localization of known and newly identified CP proteins. This work has established a new foundation for CP protein analysis for future studies.

## Introduction

Cilia and flagella are common terms used to describe the same hair-like structure of eukaryotic cells and will therefore be used interchangeably in this paper. It is known that defective cilia are implicated in a variety of different human diseases, from developmental disorders to metabolic syndromes[1]. However not all cilia are alike. Primary cilia are nonmotile and are commonly reported as sensory receptors [2]. Motile cilia, on the other hand, show beating motion at high frequencies driven by motor protein dyneins [3]. This rudimentary motion is the driving force for a plethora of multi-level systems from single cell movement to mammalian organ function and maintenance [1]. Motile cilia present in our respiratory system beat together in order to clear mucus build up and infectious agents [4]. Cilia-related diseases, otherwise known as ciliopathies, such as primary ciliary dyskinesia (PCD) are derived from the impairment of motile cilia [5, 6]. PCD is a rare congenital disease caused by defects in motile cilia. Patients who suffer from PCD often experience a wide spectrum of symptoms ranging from male infertility to an increased susceptibility to respiratory infections [7]. Failure to recognize or diagnose PCD early on often can be lethal later in life [4]. A common practice used to diagnose PCD is a cross-section analysis of patient nasal epithelium cilia using transmission electron microscopy (TEM). Due to the diversity of PCD mutations, however, many different defective proteins can lead to similar malformations [7]. In addition to this, not all mutations produce visible differences at standard TEM resolution level while they still induce PCD like symptoms. The largest obstacle to our understanding of cilia-related defects is our limited comprehension of the proteins that make up the cilia.

The cilia are highly complex structures composed from different compartments. Motile cilia consist of nine doublet microtubules (DMTs) surrounding a pair of singlet microtubules called the central pair (CP) [8]. This specific arrangement defines the “9+2” structure of the axoneme (Fig. 1A). There exists axonemal dyneins (outer dynein arm, ODA; and inner dynein arm, IDA) attached to DMTs which are responsible for the beating of cilia. Radial spoke (RS) complexes are extending from DMTs toward the CP. Intraflagellar transport (IFT) driven by IFT dyneins and IFT kinesins takes place on the DMTs [9]. This arrangement of the axoneme structure is highly conserved in all eukaryotes with motile cilia, suggesting that there exists a similar set of proteins and processes required for similar functional output. Thus, we can study the axoneme composition using model organisms like a green alga *Chlamydomonas reinhardtii*.

**Figure 1:**
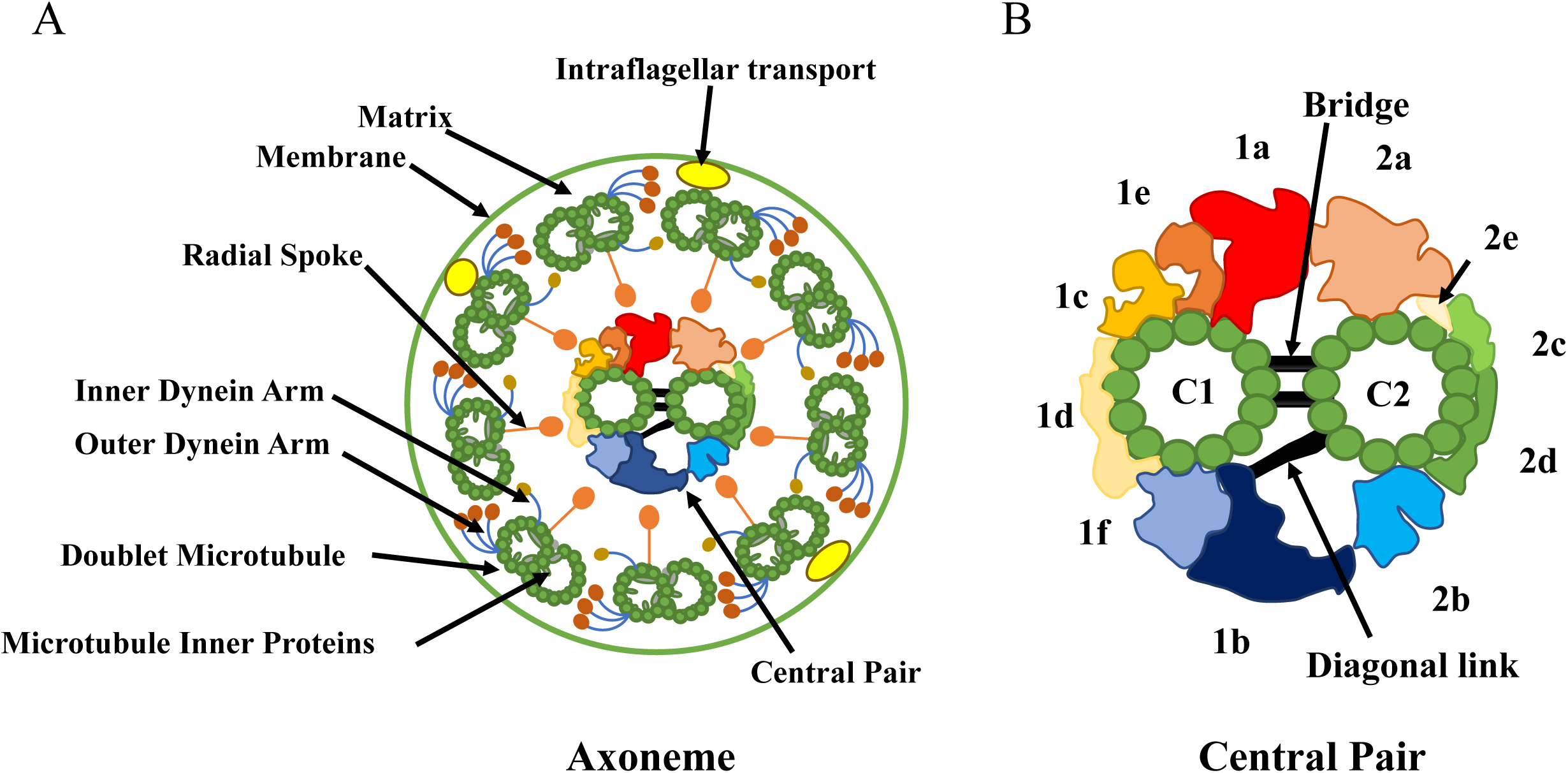
Schematic diagrams of the axonemal (A) and the CP (B) viewed from the base of flagella. The axoneme of cilia and flagella consists of nine DMTs radially surrounding the CP complex. The DMTs are decorated with ODA and IDA complexes and RS complexes. IFT takes place at the space between the membrane and the DMTs. The CP consists of two structurally dimorphic singlets termed as the C1 and C2 connected by the bridge. Several distinct sub-structures bind around the singlets with a repeating pattern along the axis of the axoneme. Diagonal link is also known to connect the C2 with the C1b region. The model of CP structure is adopted from [15].

The presence of the CP distinguishes motile cilia from its immotile counterpart, primary cilia. The CP works in a diverse array of function including the regulation of local Ca^2+^ concentration, ATP/ADP concentration and axonemal dynein activities through mechanical interactions with RS[10–13]. The CP is a huge protein complex composed of a pair of structurally and functionally dimorphic singlet microtubules named C1 and C2 and many other associated proteins [14]. The CP has a variety of sub-structures such as C1a, C1b, C1c and C1d on the C1 singlet, or C2a, C2b and C2c on the C2 singlet as characterized by traditional cross-sectional electron microscopy (EM) (Fig. 1B). C1 and C2 microtubules are connected by a structure called the “bridge” and “diagonal link”. With recent higher resolution cryo-electron tomography (cryo-ET) structures, more details of these sub-structures have been characterized allowing the C1a to be sub-classified as C1a/e, C1b as C1b/f, C2a as C2a/e and C2c as C2c/d[15]. Through this manuscript, we follow the newer nomenclature of the CP sub-structures as in Fig. 1B. These sub-structures bind with 16- or 32-nm repeating units along the axonemal axis[15]. Despite its unique existence in motile cilia and its importance to motility, the proteins that comprise the CP remain mostly unknown. Traditionally, 22 proteins (apart from α- and β-tubulins) have been characterized as components of CP (Table 1). For example, kinesin-like protein 1 (KLP1) is a phosphoprotein that localizes at the C2 microtubule around C2c/d region[16]. Mutations in several known CP proteins such as Hydin located at the C2b/f, and FAP221 (PCDP1) at the C1d have been previously shown to cause ciliopathic symptoms [10, 17]. However, it is generally believed that there should be more unidentified proteins inside the CP complex. For instance, CP-specific kinesin other than KLP1 was detected by Western blots, without knowledge about its identity[18]. Due to random insertions of transgenes into *Chlamydomonas reinhardtii* genome, previous characterization of CP proteins largely relied on phenotype-based screening of obtained CP protein mutants[19, 20]. This approach, however, remains inefficient and biased towards proteins which produce visible phenotypes.

**Table 1.** Summary of MS results of traditionally known CP proteins.

| Names | Uniprot ID | WT<br>Exclusive unique<br>peptide count<br>(Quantitative<br>values after<br>normalization) | *pf15*<br>Exclusive<br>unique<br>peptide count<br>(Quantitative<br>values after<br>normalization) | *pf15*/WT<br>ratio (%)<br>(Quantitative<br>values were<br>used) | *p*-values<br>(WT vs *pf15*) | Localizations | References |
| --- | --- | --- | --- | --- | --- | --- | --- |
| Hydin | A8HQ52 | 68, 87, 27<br>(68, 75, 53) | 0, 0, 0<br>(0, 0, 0) | 0.0 | 0.00059 | C2b | [17, 28] |
| FAP42<br>(C1b-350) | A8J614 | 63, 59, 36<br>(68, 50, 69) | 2, 2, 2<br>(2, 2, 2) | 3.6 | 0.00062 | C1b/f | [36] |
| PF6 | Q9ATK5 | 53, 65, 32<br>(65, 72, 82) | 4, 3, 5<br>(5, 3, 5) | 6.0 | 0.00016 | C1a/e | [32, 41] |
| CPC1 | Q6J4H1 | 55, 57, 30<br>(76, 66, 74) | 6, 7, 4<br>(7, 8, 4) | 8.8 | < 0.00010 | C1b/f | [32] |
| FAP54 | A8J666 | 46, 61, 27<br>(44, 48, 49) | 0, 0, 1<br>(0, 0, 1) | 0.7 | < 0.00010 | C1d | [10, 27] |
| FAP46 | A8ICS9 | 40, 50, 35<br>(52, 44, 78) | 1, 0, 2<br>(1, 0, 3) | 2.5 | 0.0052 | C1d | [10, 27] |
| FAP74 | D4P3R7 | 32, 42, 15<br>(34, 32, 33) | 0, 0, 1<br>(0, 0, 1) | 1.0 | < 0.00010 | C1d | [10, 27] |
| FAP69<br>(C1b-135) | A8IF19 | 16, 24, 11<br>(17, 23, 26) | 1, 2, 2<br>(1, 2, 2) | 8.3 | 0.0023 | C1b/f | [36] |
| PF16 | A8J0A5 | 17, 19, 16<br>(34, 69, 74) | 1, 1, 1<br>(1, 1, 1) | 1.9 | 0.010 | C1 | [22] |
| HSP70* | A8JEU4 | 14, 22, 6<br>(17, 18, 10) | 5, 5, 2<br>(7, 6, 2) | 33 | 0.028 | C1b/f | [36, 38] |
| KLP1 | A8I9T2 | 15, 16, 3<br>(17, 13, 7) | 1, 2, 1<br>(1, 2, 2) | 15 | 0.025 | C2c/d | [16, 42] |
| FAP101 | A8I345 | 13, 16, 14<br>(17, 20, 30) | 0, 0, 1<br>(0, 0, 1) | 1.6 | 0.0055 | C1a/e | [32] |
| Enolase* | A8JH98 | 17, 12, 9<br>(23, 31, 17) | 7, 9, 6<br>(9, 11, 8) | 39 | 0.017 | C1b/f | [36] |
| FAP221<br>(Pcdp1) | A8J6X7 | 7, 9, 7<br>(6, 5, 12) | 0, 0, 0<br>(0, 0, 0) | 0.0 | 0.015 | C1d | [10, 27] |
| FAP114<br>(C1a-32) | Q45QX5 | 7, 7, 5<br>(13, 8, 16) | 0, 0, 0<br>(0, 0, 0) | 0.0 | 0.0081 | C1a/e | [32] |
| FAP119<br>(C1a-34) | Q45QX4 | 7, 7, 3<br>(8, 7, 5) | 0, 0, 1<br>(0, 0, 1) | 5.1 | 0.0034 | C1a/e | [32] |
| FAP297<br>(WDR93) | A8HQE0 | 4, 8, 2<br>(6, 6, 3) | 0, 0, 0<br>(0, 0, 0) | 0.0 | 0.0040 | C1d | [27] |
| PF20 | A8ITB4 | 3, 7, 2<br>(3, 6, 3) | 1, 0, 0<br>(1, 0, 0) | 9.9 | 0.028 | C1-C2 bridge | [39] |
| PP1a* | Q9XGU3 | 2, 2, 1<br>(2, 2, 2) | 0, 0, 0<br>(0, 0, 0) | 0.0 | < 0.00010 | C1 | [43] |
| FAP174 | A8I439 | 1, 2, 4<br>(2, 4, 13) | 3, 2, 4<br>(5, 2, 8) | 79 | 0.75 | C2<br>C1b/f | [44]<br>[37] |
| Calmodulin* | A8IDP6 | 1, 3, 3<br>(1, 2, 7) | 4, 4, 4<br>(6, 5, 5) | 160 | 0.34 | C1a/e | [31, 32] |
| FAP227<br>(C1a-18) | Q45QX6 | n.d. | n.d. | - | - | C1a/e | [32] |

Three values from biological triplicate for both exclusive unique peptide count and quantitative values are shown.
Asterisks denote that these proteins are shared with other compartments of the axoneme.

In this study, we used a more comprehensive approach taking advantage of relative quantitative mass spectrometry (MS) comparing *Chlamydomonas* strains with and without intact CP. By doing so, we identified 37 new CP protein candidates. Using several different *Chlamydomonas* mutant strains of CP, we also localized these new CP proteins to certain CP sub-structures. Our results have established a new foundation for understanding the CP architecture.

## Materials and Methods

### Strains and culture condition

*Chlamydomonas reinhardtii* strains used in this study are as follows: CC-124 (Wild Type, WT), CC-1033 (*pf15*, central pair-less) [21], CC-5148 (*cpc1*, C1b/f mutant) [14], CC-1034 (*pf16*, C1 unstable) [22] and CC-1029 (*pf6*, C1a/e mutant) [23]. The cells were purchased from *Chlamydomonas* resource center and cultured in Tris-acetatephosphate (TAP) liquid media[24] with shaking or stirring, or on TAP solid plate containing 1.5% agar, on a 12 h light and 12 h dark cycle. For flagella purification, each *Chlamydomonas* strain was cultured in 1 L liquid TAP media with stirring until OD_600_ reached around 0.5-0.6.

### *Chlamydomonas* flagella isolation and purification of microtubule fraction

The cells were harvested by low-speed centrifugation (700g for 7 min at 4℃), and flagella were removed from the cell bodies by pH shock[25]. Cell bodies were removed by low speed centrifugation (1,800g for 5 min at 4℃) in HMDS (10mM HEPES, pH7.4, 5mM MgSO4, 1mM DTT, 4% sucrose containing 10 μg/ml aprotinin and 5 μg/ml leupeptin) and flagella were collected by higher-speed centrifugation (4,696g for 40 min at 4℃). Isolated flagella were resuspended in HMDEKP buffer (30 mM HEPES, pH 7.4, 5 mM MgSO4, 1 mM DTT, 0.5 mM EGTA, 25 mM potassium acetate, 0.5% polyethylene glycol, (MW 20,000) containing 10 μM paclitaxel, 1 mM Phenylmethylsulfonyl fluoride (PMSF), 10 μg/ml aprotinin and 5 μg/ml leupeptin). Paclitaxel, PMSF, leupeptin and aprotinin were added to the buffer throughout the purification after this step. Flagella were demembraned by incubating with HMDEKP buffer containing 1.5% NP40 for 30 min on ice. For cryo-electron microscopy (cryo-EM), sonication was performed for better splitting of axoneme after NP40 treatment. *Chlamydomonas* axonemes were then spun down by table top centrifuge (7,800g for 10 min at 4℃). To split the bundle of axoneme axonemes were incubated with final 1 mM ADP for 10 min at room temperature to activate dynein and then incubated with 0.1 mM ATP for 10 min at room temperature to induce doublet sliding and spun down (16,000g for 10 min at 4℃). Protease was not added for splitting. After this, *Chlamydomonas* microtubule fraction was incubated twice with HMDEKP buffer containing 0.6 M NaCl for 30 min on ice, spun down (16,000g for 10 min at 4℃), and resuspended in HMDEKP buffer. Purification process was performed three times for each strain for biological triplicates.

### Cryo-electron microscopy

3.5 μl of microtubule fraction sample (∼4 mg/ml) purified from WT *Chlamydomonas* was applied to glow-discharged holey carbon grids (Quantifoil R2/2), blotted and vitrified in liquid ethane using the Vitrobot Mark IV (FEI Company). Micrographs were obtained at 59kx nominal magnification on the direct electron detector Falcon II with the FEI Titan Krios using a total dose of ∼28 electrons/Å^2^ and 7 frames (calibrated pixel size of 1.375 Å/pixel). The defocus range was between −1.2 and −3.8 μm.

### Whole gel MS analysis

4x Laemmli buffer (#1610747, Bio-Rad) was added to the microtubule fraction samples in HMDEKP buffer so that it will be 1x, and 25-30 µg protein was loaded on the SDS-PAGE gel. Electrophoresis was performed, but the run was terminated before the proteins entered the separation gel. A band containing all proteins in the sample was then cut out from the gel and subjected to in-gel digestion[26]. Obtained peptides (∼2 μg) were chromatographically separated on a Dionex Ultimate 3000 UHPLC. First, peptides were loaded onto a Thermo Acclaim Pepmap (Thermo, 75 µm ID × 2 cm with 3 µm C18 beads) precolumn, and then onto an Acclaim Pepmap Easyspray (Thermo, 75 µm × 25 cm with 2 µm C18 beads) analytical column and separated with a flow rate of 200 nl/min with a gradient of 2-35% solvent (acetonitrile containing 0.1% formic acid) over 2 hours. Peptides of charge 2+ or higher were recorded using a Thermo Orbitrap Fusion mass spectrometer operating at 120,000 resolution (FWHM in MS1, 15,000 for MS/MS). The data was searched against *Chlamydomonas reinhardtii* protein dataset from UniProt (https://www.uniprot.org/).

### Data analysis

MS data were analyzed by Scaffold_4.8.4 (Proteome Software Inc.). Proteins with mean values of exclusive unique peptide count of 2 or more in the WT MS results were used for analysis. Raw MS data were normalized by total spectra. To identify CP protein candidates, Student’s *t*-test was applied to *pf15* and WT MS results using biological triplicates. Proteins exhibited a minimum four-fold change and the statistical significance threshold (*p* < 0.05) in *pf15* result, or proteins which were completely missing in *pf15* result were identified as new CP candidates. For statistical analysis using several mutant strain MS results, one-way analysis of variance (ANOVA) followed by Dunnett’s multiple comparisons test was performed by GraphPad Prism 8.

## Results and Discussion

### Purification of the axoneme fraction retaining the CP proteins with reduced amounts of unrelated proteins

Previous approach targeting each CP protein one by one was a time-consuming process [10, 14, 17, 18, 21–23, 27, 28]. Here, we aimed to obtain the whole CP proteome using MS. Due to the sensitive nature of MS, peptide detection tends to have an unfavorable preference for large and abundant proteins. In our previous paper, we used a ciliate *Tetrahymena thermophila* as a model organism[29], but the green alga *Chlamydomonas reinhardtii* was used in this study since there were contaminated mucocyst proteins in the MS of microtubule fraction purified from *Tetrahymena* (unpublished data). Previous proteomic analysis of whole *Chlamydomonas* flagella also showed the presence of an abundant amount of proteins from membrane and matrix fractions, and large proteins such as dynein heavy chains [30]. Thus, it was important to prepare samples for MS with reduced amount of unrelated proteins which would hinder the detection of CP proteins, while the CP structure with its associated proteins remains undisturbed. To achieve this, microtubule fraction was purified from WT *Chlamydomonas* flagella by sequential purification following the deflagellation by pH shock (Fig. 2A). First, proteins from the membrane and matrix fraction were removed by NP-40 treatment. Demembranated axonemes were treated with 0.6 M NaCl twice to extract axonemal dyneins as much as possible. From SDS-PAGE, significant amounts of proteins were removed in the final extract leaving the tubulin band which is a main component of CP and DMT (Fig. 2B). Though we also tried to remove RS complexes by dialysis against low salt buffer[29] or KI treatment[31], it was not possible to remove *Chlamydomonas* RS complex keeping microtubule structures unaffected. Thus, the RS complexes were left in our sample. To detect all proteins in the sample, the purified microtubule fraction was analyzed by whole gel MS (Materials and Methods) and almost all (21 out of 22) known CP proteins such as Hydin, CPC1, Pf6, FAP69, Pf16, KLP1 and FAP221 (PCDP1) were detected (Table 1). The only known CP protein which we failed to detect was FAP227 (C1a-18) [32]. Since the size of FAP227 is small (18 kDa), it is thought to be unfavorable for the detection by MS. Known CP proteins detected were previously shown to localize at different CP sub-structures (Table 1). Microtubule inner protein (MIP) candidates like Rib43a, Rib72 and Tektin[33], and RS proteins were also detected since these structures are tightly associated with DMTs (Supplementary Excel File 1). Proteins from other axonemal components such as IFT complex proteins, IFT dyneins, IFT kinesins, axonemal dyneins, and dynein regulatory complex were still detected due to the high sensitivity of MS though we tried to reduce them as much as possible. Twice salt treated samples were imaged using cryo-EM and singlet microtubules from CP with characteristic repeating protrusions were observed along with DMTs (Fig. 2C and Fig. S1). Together with this cryo-EM result, we concluded that our purification method retained the CP proteins and was usable for proteomic analysis.

**Figure 2:**
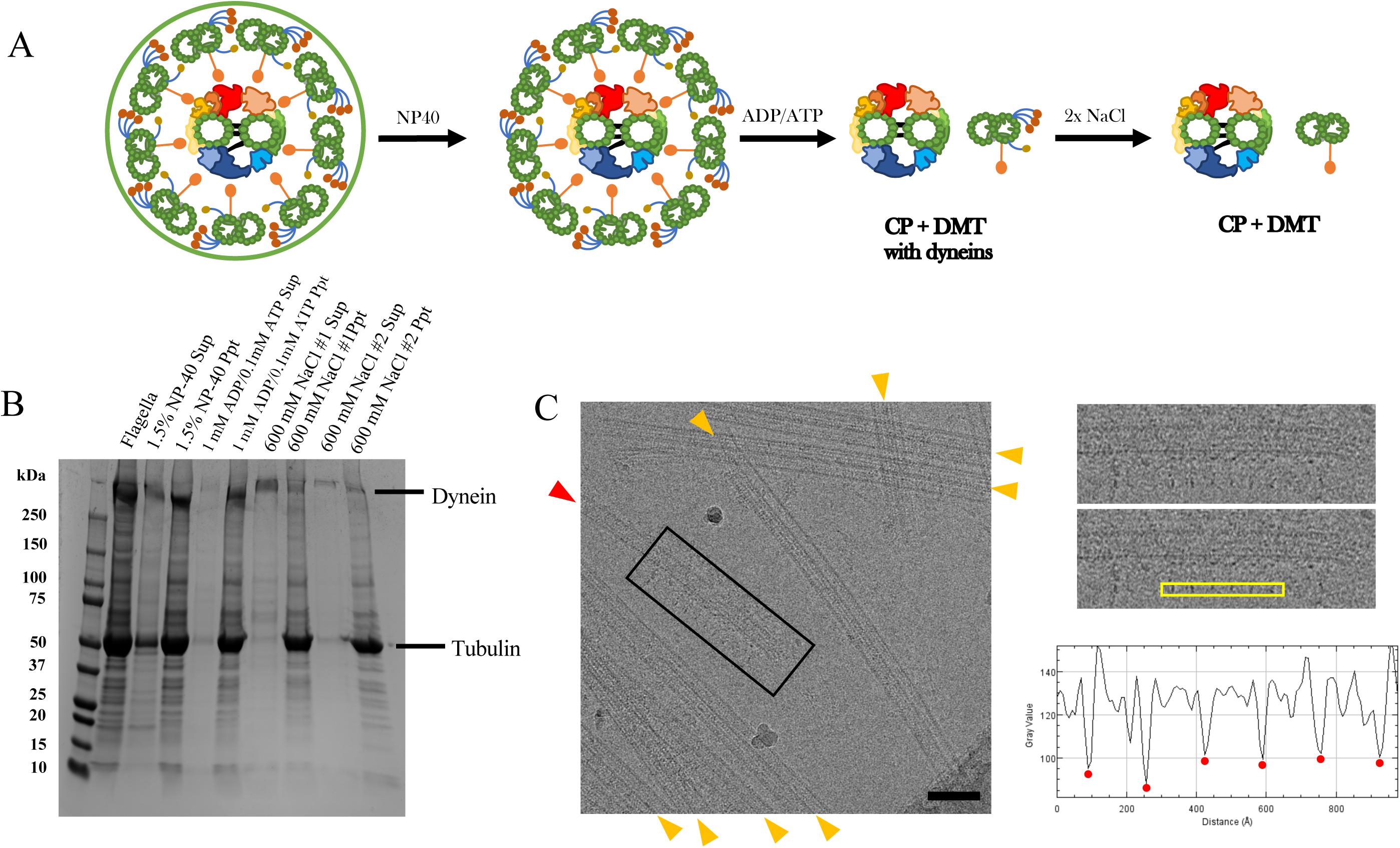
Preparation of microtubule fraction for MS. (A) A schematic diagram of sequential purification of the axoneme. Flagella were demembraned using a detergent NP-40 following isolation from *Chlamydomonas* cells. Demembranated axoneme was incubated with ADP and ATP to induce splitting of the DMTs and the CP and then treated with 0.6 M NaCl twice to shed large protein complexes such as dyneins. (B) SDS-PAGE gel demonstrating protein shedding after sequential purification. The signal of dynein heavy chain band (> 500 kDa) was decreased significantly after NaCl treatments. In contrast, the tubulin band which is a main component of the CP and DMTs showed little change after sequential purification. (C) A typical cryo-EM image of purified sample showing the presence of singlet microtubule from the CP. In our cryo-electron micrographs of our purified microtubule fraction, both DMTs (orange arrowheads) and singlet microtubule from the CP (red arrowhead) with characteristic protruding sub-structures were observed (see also Fig. S1). Boxed area of the micrograph is shown in the right panel. The plot profile of yellow box area was generated by ImageJ and the distances between the peaks (red dots) were measured. The averaged distance between the protrusions was 16.7 nm which is consistent with the known repeating unit of the CP[15]. There were more numbers of the DMTs compared with singlets from the CP reflecting the stoichiometry inside the axoneme. Scale bar represents 100 nm.

### Identification of new CP proteins by comparative proteomic analysis

Next, we sought to characterize new CP protein candidates. Since it is not possible to sub-fractionate CPs from DMTs as they are both microtubule-based structures, we decided to use a comparative proteomic approach using a specific *Chlamydomonas* mutant lacking whole CP complex. *Chlamydomonas* mutant strain *pf15* contains a mutation in a gene encoding p80 subunit of microtubule severing enzyme Katanin[21]. The resulting effects prevent the entire CP complex from assembling and lead to paralyzed flagella while leaving other components like DMTs intact (Fig. 3A). *pf15* strain was chosen among other CP-less mutants in this study since it was previously shown that the flagella length of *pf15* strain is closer to that of WT compared with other CP-less mutants[34]. An identical purification process as WT was used for *pf15* mutant flagella, and the MS analysis was performed from the microtubule fraction of *pf15* (Fig. 3A and B). Similar amount of *pf15* microtubule fraction as WT was analyzed by whole gel MS, and these results were normalized by total spectra and compared (see details for Materials and Methods). In the MS result of *pf15* sample, there were many proteins significantly reduced or completely missing compared with the WT result (Fig. 3C). Proteins known to be tightly associated with DMTs including Rib43a, Rib72 and Tektin and RS proteins were detected at the similar levels with WT (Supplementary Excel File 2). In contrast, 18 out of 20 known CP proteins detected in WT were completely missing or significantly decreased in *pf15* result (Table 1). Calmodulin did not show significant decrease in the *pf15* MS result. Since Calmodulin is shared with the RS[31] the decrease of Calmodulin is thought to be masked by signals from the remaining RS. FAP174 also did not show significant decrease but this could mean that FAP174 is also present in other axonemal structures as well (discussed later).

**Figure 3:**
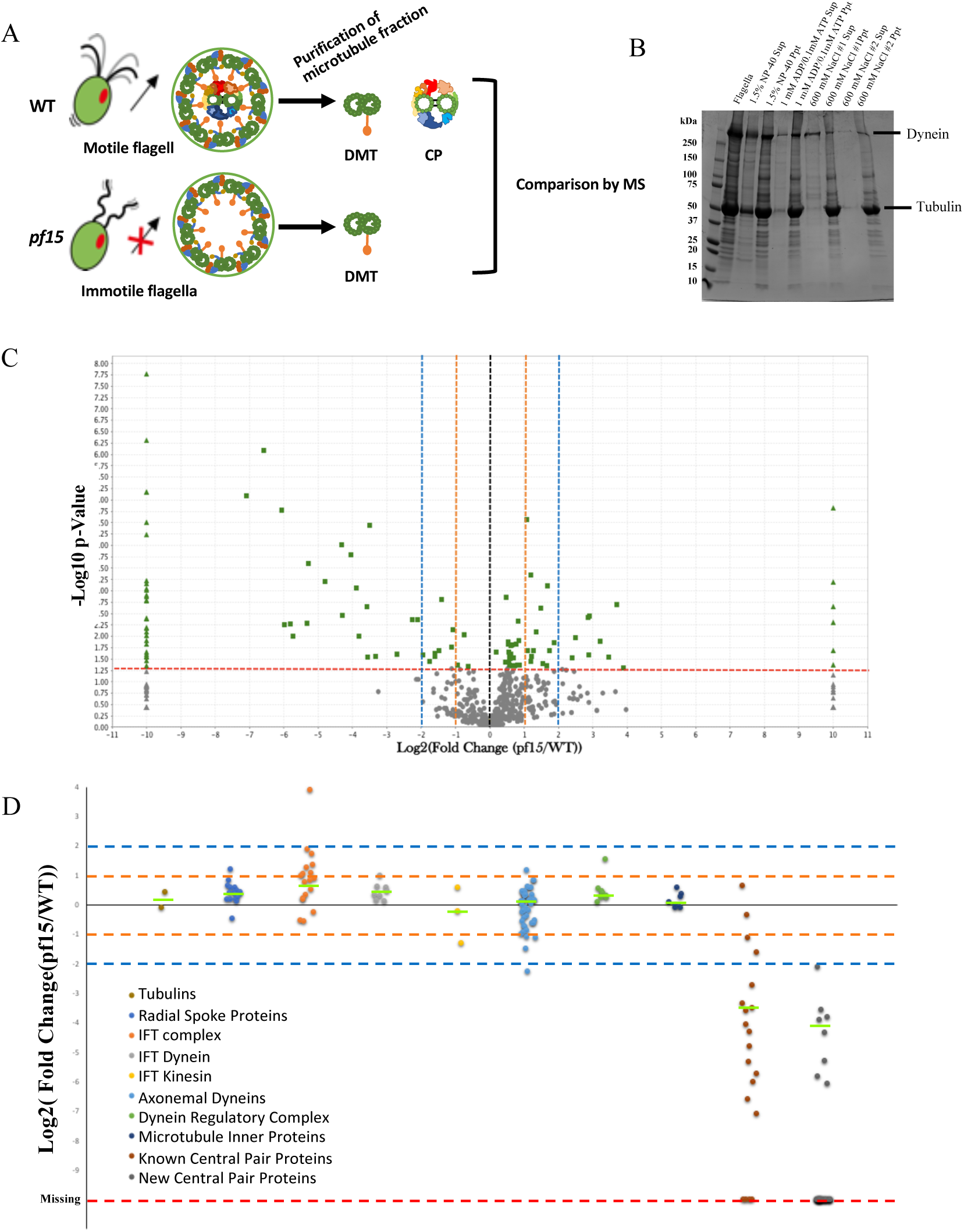
Identification of new CP proteins by MS. (A) Schematic diagrams of the axoneme structures from WT and *pf15 Chlamydomonas* strains and expected microtubule structures obtained from either axoneme structure. The DMTs and CP complexes were purified from WT flagella while only DMTs are expected to be purified from *pf15* flagella. Obtained microtubule fractions from WT and *pf15* were analyzed by MS and the results were compared. (B) SDS-PAGE result of sequential purification of microtubule fraction from *pf15* flagella showing similar pattern with that of WT flagella. (C) Volcano plot of comparison of WT and *pf15* mass spectrometry results. Changes in a protein abundance between WT (n = 3) and *pf15* (n = 3) results were plotted on a volcano plot. Dashed red line indicates the significance threshold of *p* < 0.05 and proteins meet this criterion are shown in green. Triangle dots represent completely missing proteins in either WT or *pf15* result. Two-and four-fold changes are shown by the orange and blue dashed lines, respectively. There were more proteins completely missing in *pf15* results while many others showed more than two-fold decrease in *pf15* results. (D) Plot of fold changes of proteins categorized into different groups. Proteins identified by MS were arranged by groups (Tubulins; RS proteins; IFT complex proteins; IFT dynein; IFT kinesin; axonemal dyneins; dynein regulatory complex; MIP candidates; known CP proteins) and fold change between WT and *pf15* results of each protein was plotted. Two-and four-fold changes are shown by the orange and blue dashed lines, respectively. Green lines indicate the median value for each category. Statistical significance compared with tubulin result was examined by ANOVA followed by Dunnett’s multiple comparisons test. Among these classes, only known CP protein class was significantly reduced with *p*-value of 0.0005. Fold changes of our new CP protein candidates are also shown at the rightmost column. Red line represents proteins that were completely missing in *pf15*. Proteins included in each class are listed in Tables 1 and 2, and Supplementary Excel File 2.

**Table 2.** MS results of new CP proteins and their localizations inside the CP complex.

| Names | Uniprot ID | WT exclusive unique peptide counts (quantitative values) | *pf15* exclusive unique peptide counts (quantitative values) | *pf15*/WT ratio (%) (quantitative values were used) | *p*-values (WT vs *pf15*) | Localizations |
| --- | --- | --- | --- | --- | --- | --- |
| CHLREDRAFT_150638 | A8J566 | 1, 5, 0 (1, 4, 0) | 0, 0, 0 (0, 0, 0) | 0 | 0.23 | C1a/e (this study) |
| CHLREDRAFT_170023 | A8IMQ8 | 3, 9, 0 (3, 6, 0) | 0, 0, 0 (0, 0, 0) | 0 | 0.17 | C1b/f (this study) |
| CHLREDRAFT_177061 | A8J9A4 | 7, 7, 2 (6, 5, 3) | 0, 0, 1 (0, 0, 1) | 7.1 | 0.0098 | C1b/f (this study) |
| DPY30** | A8J1X7 | 1, 2, 1 (1, 1, 2) | 0, 0, 0 (0, 0, 0) | 0.0 | 0.0041 | C1a/e ([37] & this study) |
| EF-3 | A8ISZ1 | 4, 2, 1 (4, 1, 2) | 0, 0, 0 (0, 0, 0) | 0.0 | 0.047 | Not assigned |
| FAP7* | A8IVW2 | 14, 16, 7 (26, 33, 28) | 1, 0, 1 (1, 0, 1) | 2.6 | 0.00025 | C1a/e (this study) |
| FAP47** | A8IPW8 | 22, 35, 15 (23, 27, 26) | 0, 0, 0 (0, 0, 0) | 0.0 | < 0.00010 | C2b (this study) |
| FAP65* | A8JFU2 | 13, 23, 10 (12, 19, 16) | 0, 0, 0 (0, 0, 0) | 0.0 | 0.0016 | C2a,c,d,e and Bridge (this study) |
| FAP70* | A8I7W0 | 12, 22, 11 (14, 26, 23) | 0, 1, 0 (0, 1, 0) | 33 | 0.0053 | C2a,c,d,e and Bridge (this study) |
| FAP75* | A8HYW3 | 13, 19, 8 (13, 16, 15) | 0, 1, 1 (0, 1, 1) | 5.0 | < 0.00010 | C2a,c,d,e and Bridge (this study) |
| FAP76** | A8J128 | 24, 26, 14 (26, 23, 26) | 0, 1, 0 (0, 1, 0) | 1.5 | < 0.00010 | C1d (this study) |
| FAP81* | A8IPC1 | 23, 27, 12 (24, 24, 23) | 0, 0, 0 (0, 0, 0) | 0.0 | < 0.00010 | C1d (this study) |
| FAP92* | A8HR45 | 28, 30, 17 (29, 23, 36) | 0, 0, 0 (0, 0, 0) | 0.0 | 0.0017 | C1d (this study) |
| FAP99** | A8IUG5 | 9, 13, 1 (10, 9, 2) | 0, 0, 0 (0, 0, 0) | 0.0 | 0.060 | C1 [37]<br>C2a,c,d,e and Bridge (this study) |
| FAP105* | A8IKV8 | 3, 5, 0 (3, 4, 0) | 0, 0, 0 (0, 0, 0) | 0.0 | 0.12 | C1d (this study) |
| FAP108* | A8IPA9 | 2, 3, 1 (2, 2, 2) | 0, 0, 0 (0, 0, 0) | 0.0 | 0.00090 | C1d (this study) |
| FAP123* | A8IEJ6 | 4, 3, 0 (4, 2, 0) | 0, 0, 0 (0, 0, 0) | 0.0 | 0.16 | C2a,c,d,e and Bridge (this study) |
| FAP125* | A8IY87 | 14, 18, 9 (15, 18, 16) | 0, 0, 0 (0, 0, 0) | 0.0 | < 0.00010 | C1c (this study) |
| FAP147* | A8IT32 | 7, 10, 3 (6, 7, 7) | 0, 0, 0 (0, 0, 0) | 0.0 | < 0.00010 | C2a,c,d,e and Bridge (this study) |
| FAP171* | A8IUF4 | 4, 9, 2 (4, 7, 3) | 1, 0, 0 (1, 0, 0) | 8.4 | 0.029 | C2a,c,d,e and Bridge (this study) |
| FAP173 | A8JAF7 | 3, 3, 1 (4, 2, 2) | 0, 0, 0 (0, 0, 0) | 0.0 | 0.012 | Not assigned |
| FAP194* | A8J5U4 | 9, 13, 4 (10, 10, 7) | 0, 0, 0 (0, 0, 0) | 0.0 | 0.0013 | C2a,c,d,e and Bridge (this study) |
| FAP199 | A8J1E6 | 1, 2, 3 (1, 1, 7) | 0, 0, 0 (0, 0, 0) | 0.0 | 0.19 | Not assigned |
| FAP209 | A8J100 | 6, 8, 3 (6, 5, 5) | 0, 1, 0 (0, 1, 0) | 6.7 | 0.00086 | C1c (this study) |
| FAP216* | A8JGM3 | 12, 16, 4 (13, 13, 7) | 0, 0, 0 (0, 0, 0) | 0.0 | 0.0064 | C1e to f? (this study) |
| FAP219 | A8J9I0 | 5, 7, 1 (5, 5, 2) | 0, 0, 0 (0, 0, 0) | 0.0 | 0.022 | C1c (this study) |
| FAP225* | A8HNF2 | 14, 19, 4 (14, 22, 7) | 0, 0, 0 (0, 0, 0) | 0.0 | 0.034 | C2a,c,d,e and Bridge (this study) |
| FAP239* | A8J319 | 0, 5, 2 (0, 3, 3) | 0, 0, 0 (0, 0, 0) | 0.0 | 0.12 | C2a,c,d,e and Bridge (this study) |
| FAP244 | A8IZG0 | 12, 14, 5 (14, 10, 12) | 0, 0, 0 (0, 0, 0) | 0.0 | 0.00069 | C2a,c,d,e and Bridge (this study) |
| FAP246** | A8HNZ7 | 7, 6, 3 (7, 6, 5) | 0, 0, 0 (0, 0, 0) | 0.0 | 0.00069 | C1b/f ([37] & this study) |
| FAP266* | A8JB69 | 4, 5, 2 (4, 3, 3) | 1, 1, 0 (1, 1, 0) | 23 | 0.0043 | C2a,c,d,e and Bridge (this study) |
| FAP279 | A8HWC6 | 5, 7, 1 (6, 4, 2) | 0, 0, 0 (0, 0, 0) | 0.0 | 0.029 | C1d (this study) |
| FAP289* | A8JCZ9 | 8, 12, 4 (9, 9, 8) | 0, 0, 0 (0, 0, 0) | 0.0 | < 0.00010 | C1d (this study) |
| FAP312* | A8IUV6 | 2, 5, 1 (2, 3, 2) | 0, 0, 0 (0, 0, 0) | 0.0 | 0.0066 | Not assigned |
| FAP348* | A8JBI2 | 2, 3, 2 (2, 2, 3) | 0, 0, 0 (0, 0, 0) | 0.0 | 0.0095 | C1a/e (this study) |
| FAP412* | A8JGL8 | 6, 9, 0 (6, 6, 0) | 0, 0, 0 (0, 0, 0) | 0.0 | 0.12 | Not assigned |
| MOT17* | A8J798 | 3, 4, 3 (3, 2, 7) | 0, 0, 0 (0, 0, 0) | 0.0 | 0.044 | C1c (this study) |
| Phosphoglycerate mutase | A8HVU5 | 4, 10, 0 (4, 7, 0) | 0, 0, 0 (0, 0, 0) | 0.0 | 0.15 | C1b/f? (this study) |

\*: CP candidates shared with [37].
\*\* : CP proteins which were confirmed by [37].
DPY30 was not originally in our CP candidates but shown here since it was confirmed as CP protein.

To clearly see the differences between WT and *pf15* results, we performed comparison based on protein categories which include, tubulins, IFT complex, IFT-dynein, IFT-kinesin, RS proteins, axonemal dyneins, dynein regulatory complex proteins, MIPs, and known CP proteins (Fig. 2B). Since values were normalized, α- and β-tubulins were detected at the same level between WT and *pf15* results and served as a control (Fig. 3D). As clusters, only known CP proteins were significantly decreased in *pf15* MS results, validating our strategy. In our *pf15* MS result, IFT complex proteins were slightly up-regulated similarly with previous observations[34] but not significantly in our condition. Other classes like IFT dynein, IFT kinesin, axonemal dyneins, dynein regulatory complex proteins, and MIPs did not show any significant decrease as compared to known CP proteins category. From these results, we concluded that our method is valid to identify CP proteins.

Among previously characterized CP proteins was a population of previously uncharacterized proteins totally missing or significantly reduced in *pf15* result. These characterized proteins were thought to be new candidates of the CP. Though CP proteins like enolase and HSP70, which are known to be shared with other axonemal component, showed two- to four-fold decrease in *pf15* result, this area also contained proteins such as DHC9, p38 and KAP from other categories (Fig. 3D). Almost all the proteins from other complexes did not show decrease with four-fold or larger except DHC8 (Supplementary Excel File 2). Thus, proteins which were decreased significantly (p < 0.05) with a 4-fold decrease or greater in *pf15* compared with WT result, or completely missing in *pf15* were categorized as new CP proteins. 37 proteins including FAP7, FAP47, FAP65, FAP70 were identified as new CP candidates with this criterion (Fig. 3D and Table 2). One of these newly identified CP protein candidates was FAP125. FAP125 is a kinesin-like protein which was previously proposed to be another kinesin-like protein in CP without direct evidence[35]. Our study presented direct evidence showing that FAP125 is actually a novel CP-associated kinesin in addition to KLP-1. Furthermore, phosphoglycerate mutase was detected in WT result but completely missing in *pf15* result (Table 2). Phosphoglycerate mutase was previously shown to be present in the axoneme and play roles in ATP production together with enolase[36]. Interestingly, enolase which is involved in the same ATP synthesis pathway together with phosphoglycerate mutase is known to be a component of CP as well as present in the membrane and matrix fractions. Since these enzymes work together, it is likely that phosphoglycerate mutase is also integrated into the CP complex to facilitate the reaction.

During the time we were finishing this manuscript, a similar proteomic study aiming to identify CP proteins was published[37]. In this study, *pf18 Chlamydomonas* strain, which also lacks CP complex, was used to compare with WT strain instead of *pf15* used in our study. In addition, only demembranated whole axoneme structure was analyzed by quantitative MS instead of purified microtubule fractions. We also performed biological triplicate through our work to identify only consistent candidates unlike replicate in theirs. Despite the differences in strains and methods used, 26 out of 37 identified proteins in our work were shared with their results, making these proteins very strong candidates of CP proteins (Table 2). 11 proteins were assigned as new CP candidates only in our work and 19 proteins only in [37] (Table 2 and Table S1). These differences might be due to contaminated proteins from remaining axonemal components in Zhao *et al*.’s purification method. For example, proteins like FAP39, 49, 72, 139 and 154 were characterized as new CP protein candidates in Zhao et al., (2019) [37], but these proteins were consistently detected in *pf15* in our triplicate result and less likely to be a stable components of CP complex (Table S1). Conversely, it could be because some of the CP proteins fell off in our purification method. NAP was shown to be a component of the C1a/e region by immunoprecipitation by Zhao et al., [37], but we failed to detect NAP in our MS. This could be due to its weak association of NAP to the C1a/e region. DPY30 was also identified and further confirmed to be a component of CP by immunofluorescence [37]. We detected DPY30 in our MS results, but the detected amount was little and was not in our original CP list. It is also possible that differences are due to different strains used (discussed later). Nevertheless, it is noteworthy that even using different strains and methods, many common proteins were identified as new CP protein candidates. These two studies can be used in a mutually complementary way to understand the architecture of the CP complex.

### Localizing CP candidates into sub-structures of CP complex

We further aimed to identify the localizations of these new CP proteins inside CP complex. To achieve this, we used different kinds of *Chlamydomonas* mutants lacking specific CP sub-substructures *pf*6 (C1a/e mutant) and *cpc*1 (C1b/f mutant) in addition to *pf16* (unstable C1 microtubule) (Fig. 4A). Following the same sample preparations and MS conditions as WT (Fig. S2), we compared normalized MS results from five different strains allowing us to produce MS detection profile for each protein of interests (Fig. 4B-F and Fig. S3).

**Figure 4:**
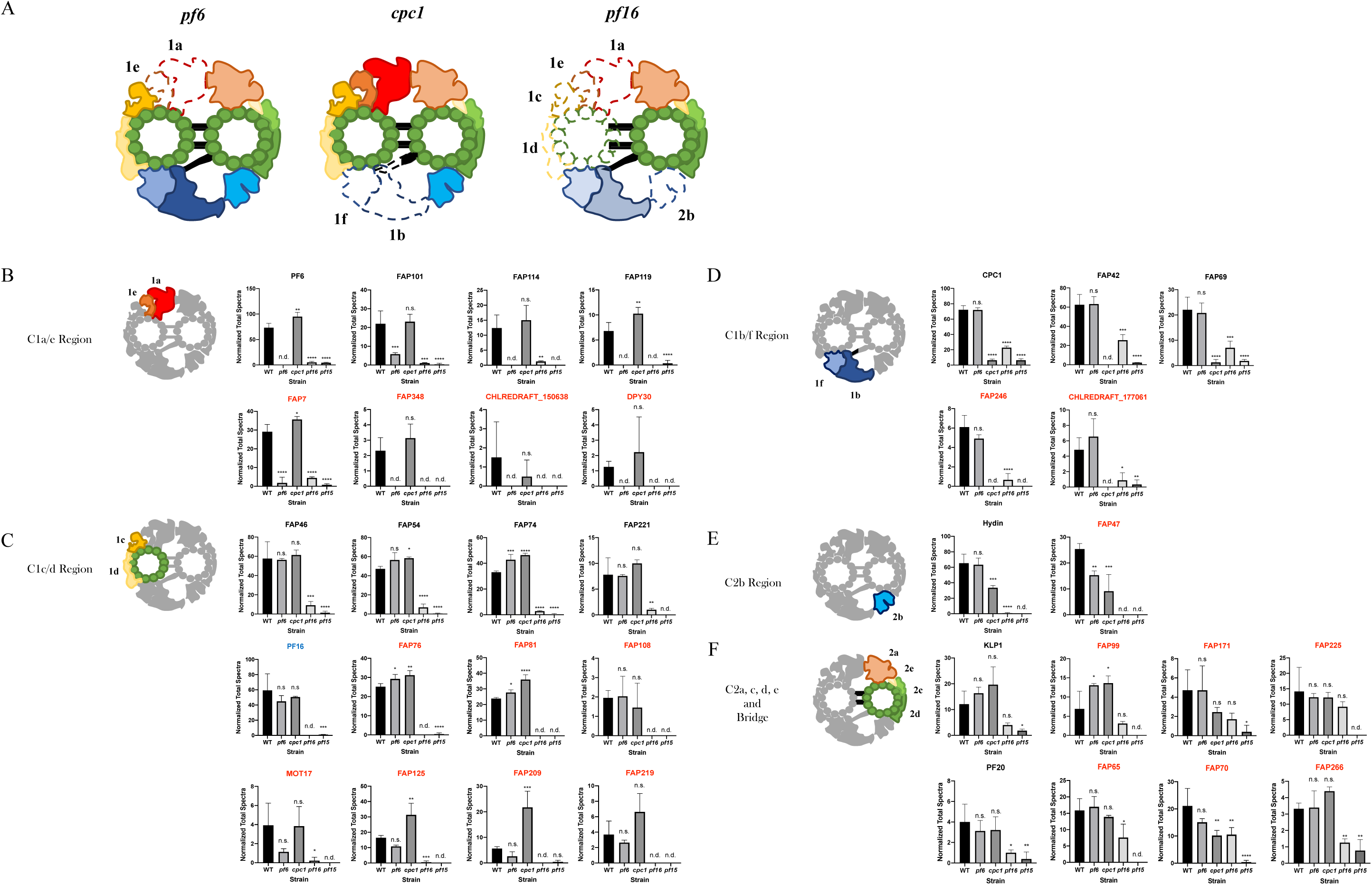
MS analyses using *Chlamydomonas* mutant strains lacking CP sub-structures. (A) Schematic diagrams of CP structures from mutants lacking sub-structures of CP. Sub-structures of CP which are missing in *pf6, cpc1* and *pf16* strains are shown in dashed lines. *pf6* strain is missing the C1a/e structure (formerly the C1a), *cpc1* strain lacks the C1b/f structure (formerly the C1b) while *pf16* strain has an unstable C1 structure. The C1b/f region is shown translucent since this region can remain attached to the C2 microtubule with the diagonal link[14]. (B-F) MS profiles of CP proteins and their possible localizations. Detected levels of proteins were compared among strains (WT, *pf6*, *cpc1*, *pf16* and *pf15*). Mean values of normalized quantitative values of each CP protein are shown (error bars represent SD for biological triplicate). Known CP proteins (black) that have been localized to specific sub-complexes showed similar MS profiles. These proteins were used as references to assign newly identified CP proteins (red) to certain sub-structures, such as the C1a/e area (B), the C1c/d area (C), the C1b/f area (D), the C2b area (E), and the C2a, c, d, e & bridge area (F). Known CP protein PF16 (blue) was also categorized into the C1c/d region based on the MS profile. Statistical test was performed by ANOVA followed by Dunnett’s multiple comparisons test comparing with WT values (\**p* < 0.05; \*\**p* < 0.01; \*\*\**p* < 0.001; \*\*\*\**p* < 0.0001; n.s., not significant; n.d., not detected). Plots not shown here are presented in Fig. S3.

Traditionally known CP proteins which localize at the same area shared similar MS profiles. CP proteins like PF6, FAP101, FAP114 (C1a-32) and FAP119 (C1a-34), which were shown to be located at the C1a/e sub-structure[23, 32], were detected both in WT and *cpc1* strains since they retain the C1a/e structure while these proteins were not detected or detected with very little amounts in *pf6*, *pf15* and *pf16* strains because of the lack of the C1a/e region (Fig. 4B). To verify this statistically, we performed ANOVA test and these known C1a/e proteins (PF6, FAP101, FAP114 (C1a-32) and FAP119 (C1a-34)) were all significantly decreased in *pf6*, *pf15* and *pf16* strains but not in *cpc1* strain (Fig. 4B). Among our new CP candidates, FAP7, FAP348 and CHLREDRAFT_150638 showed the same pattern by ANOVA test and thought to be present in the C1a/e region (Fig. 5, and Table 2). Though it was not in our original CP candidates, DPY30 which was proposed to be located at the C1a/f region by immunoprecipitation[37], also met this standard further verifying our assignment.

**Figure 5:**
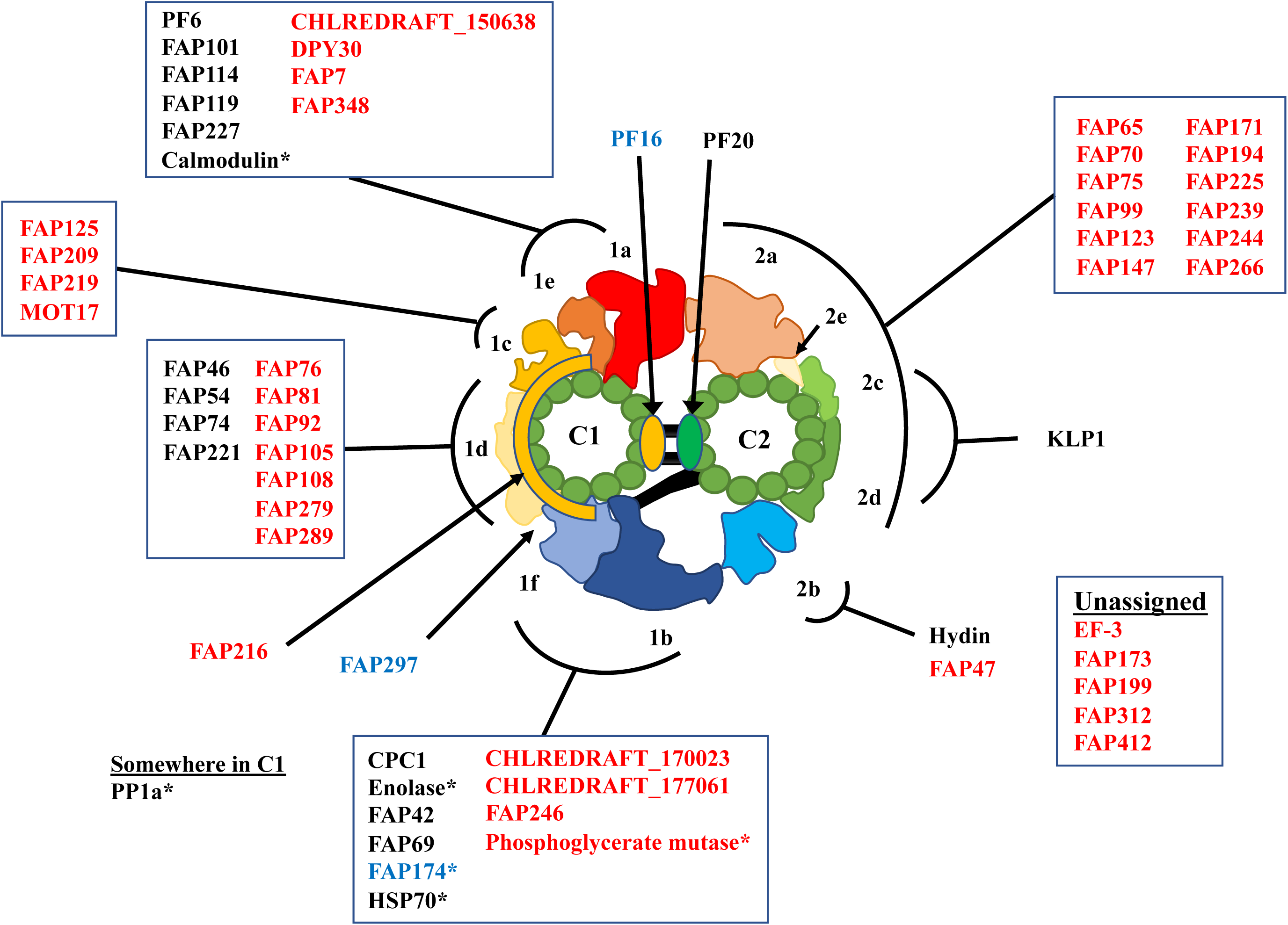
Summary of localizations of known and new CP proteins. Proteins are mapped to CP sub-structures (C1a/e; C1c; C1d; C1b/f; C2b; and C2a, c, d, e & bridge areas) based on our MS profiles. Traditionally known CP proteins are shown in black and new CP proteins are shown in red. Known CP proteins (PF16 and FAP174) which were assigned to certain sub-structures in our work are highlighted in blue. FAP297 was proposed to localize at C1d region, but it is thought to be at the interface between C1d and f from our MS result. Asterisks denote the proteins possibly shared with other axonemal structures.

There were several CP proteins known to localize at the C1d area. FAP46, FAP54, FAP74, FAP221 (PCDP1) and FAP297 are such proteins[10]. These proteins were proposed to form a complex located at the C1d region. In our MS profile, most of these proteins show a similar trend being significantly reduced only in *pf15* and *pf16* strains since they lack this region (Fig. 4C). The only exception was FAP297 which showed significant reduction in *cpc1* strain which lacked the C1b/f region (Fig. S3A). This could mean that FAP297 is located at the interface between the C1d and the C1f and interacting with proteins from the C1f (Fig. 5). Zhao *et al*., also were not able to detect FAP297 by immunoprecipitation using FAP46 as a bait though all other known C1d proteins were detected [37]. This further supports our idea that FAP297 is located away from other C1d proteins. Thus, the trend shared between FAP46, 54, 74 and 221 was used as a standard for C1d protein candidates. Our CP candidates were classified into the C1d area based on the statistical analysis results of MS profiles. FAP76, 81, 92, 105, 108, 279 and 289 showed similar trends (Fig. 4C and S3A) and were assigned into the C1d (Fig. 5 and Table 2). FAP279 is a leucine-rich repeat-containing protein that was not assigned as a CP protein before. A homologue is also present in humans (LRRC72). FAP81 was proposed to be a component of the C1a/e projection since it was detected along with other known C1a/e proteins by immunoprecipitation using DPY30 as a bait[37]. Our result using different mutant strains does not support their conclusion of localization since FAP81 is clearly present in *pf6* mutant lacking this region. Rather, FAP81 is thought to interact with DPY30 via other protein (discussed later). Our MS profile gives insights not only for new CP candidates but also for previously known CP proteins. PF16 protein which is a known C1 protein also showed similar MS profile with other C1d proteins (Fig. 4C) and thus thought to be located at this region (discussed later).

Some proteins like MOT17, FAP125, FAP209 and FAP219 showed significant reduction only in *pf15* and *pf16* strains by ANOVA test like other C1d proteins, but these proteins were slightly reduced in *pf6* strain (Fig. 4C lowest row). These MS patterns were somewhat at the middle of known C1a/e proteins and C1d proteins. Thus, these proteins are thought to be at the interface of the C1a/e and the C1d, namely the C1c area (Fig. 5). One of these proteins, MOT17 was shown to interact with known C1a/e proteins by immunoprecipitation which is neighboring region of the C1c[37]. FAP125 was proposed to be somewhere in C1 microtubule, but its localization to certain sub-structure was not achieved. Our result located FAP125 into specific sub-structure of the C1 microtubule. For FAP209&219, there were no localization information at all.

Traditionally known proteins belonging to the C1b/f sub-complex like CPC1, FAP42 (C1b-350) and FAP69 (C1b-135) also share similar patterns distinct with significant decrease in *cpc1*, *pf15* and *pf16* but with modest decreases in *pf16* (Fig. 4D). At first, we were puzzled with this result of modest decrease in *pf16* strain since *pf16* strain is generally assumed to have an unstable C1 to which the C1b/f region is attached. Therefore, we looked into the article characterizing the *pf16* mutant carefully and realized that the C1b/f part remains with the C2 microtubule due to the diagonal link connecting these structures even other sub-structures like the C1a, c, d and e, and C2b were missing[14]. This was also mentioned by the authors but has been overlooked in recent articles. Thus, we concluded that the C1b/f region is partially present in *pf16* structure in our purification condition (Fig. 4A). Enolase and HSP70A showed different MS patterns from other known C1b/f proteins (Fig. S3B), but these proteins were previously shown to be present both in CP complex and in other compartments of the axoneme, thus the differences are thought to represent the presence of these proteins in other compartments of the axoneme[36, 38]. Therefore, the MS profile shared with CPC1, FAP42 and FAP69 was used as a standard for C1b/f proteins. Among our new CP candidates, FAP246 and CHLREDRAFT_177061 showed C1b/f-like profile (Fig. 4D) and thought to be located at this region (Fig. 5 and Table 2). FAP246 was shown to interact with other C1b/f proteins by immunoprecipitations by Zhao et al., (2019) [37] and our localization is consistent with this.

Hydin is the only protein known to be associated with the C2b region[17, 28]. Based on previous cross-sectional EM result, this region solubilizes before the C1b/f region in *pf16* CP structure[14]. Consistent with this, Hydin was missing in *pf16* MS result. Hydin was also decreased in *cpc1* compared with WT. From previous studies, the C2b projection is at close proximity to the C1b sub-structure[15]. Thus, it is possible that the interactions between neighboring sub-structures C2b and C1b are tighter than previously assumed. In our MS profile, FAP47 showed similar trend with Hydin (Fig. 4E). In recent MS result, FAP47 also showed elution pattern similar with Hydin[37]. Based on these results, FAP47 was tentatively assigned to the C2b region in our model (Fig. 5 and Table 2). In Zhao et al., (2019), FAP49, 72, 154 and 416 were also identified as CP protein and proposed to form a complex with FAP47 based on immunoprecipitation result [37]. In our MS result, FAP49, 72, & 154 were detected but not categorized as new CP candidates since they did not show significant decrease in *pf15* (Table S1). Detection of these proteins were not very consistent, but they were present in all *pf15* triplicate as WT level and completely missing in all *cpc1* triplicate (Supplementary Excel File 1). The exact reason for this is not apparent, but these proteins might be loosely attached to the C1b/f region with the aid of FAP47 at the C2b rather than being stable components of CP. It was shown that the amounts of electron-dense materials around the CP complex is less in *pf18* compared with *pf15* strain[37]. Thus, these loosely anchored proteins might correspond to these electron-dense materials.

KLP1 is known to localize at the C2c/d area[16]. KLP1 was detected in most strains but in a very small amount in *pf15* strain. This is consistent with the result of previous cross-section EM showing that the C2c/d region is stably bound to the C2 microtubule in *pf16*[14] (Fig. 4A). Interestingly, PF20 protein which is known to be localized at the “bridge” connecting the C1 and C2 microtubules showed MS profile similar with that of KLP1 being detected slightly more in *pf16* strain compared with *pf15* result (Fig. 4F). By immunogold labeling EM in previous study, gold particles were found to be bound to only one of the CP singlets, presumably the C2 microtubule[39]. Our result also suggests that PF20 is associated more tightly to the C2 microtubule (Fig. 5). PF20 was previously shown to interact with PF16 protein by yeast two hybrid study[40] but the MS profile of PF20 was somewhat different from that of PF16 notably the presence in *pf16* strain. This means that though PF16 and PF20 proteins are interacting, PF16 is more tightly bound to the C1 microtubule and PF20 to the C2 microtubule. Considering that the mutation in gene encoding PF16 results in the unstable C1 microtubule, PF16 is thought to be present on the surface of the C1 microtubule facing the C2 and anchoring the C1 microtubule to the bridge (Fig. 5). Among our new CP candidates, proteins like FAP65, 70, 75, 99, 123, 147, 171, 239, 244 and 266 showed similar MS profiles with KLP1 and PF20 (Fig. 4F & S3C) and therefore thought to be present at the C2a, c, d, e and bridge region (Fig. 5 and Table 2). The C2a/e area is also included since this part is known to be stably attached to the C2 microtubule in *pf16* strain[14]. Previous cryo-ET study has shown that there are protein structures inside the C2 microtubule tubulin lattice similar to the MIPs in the DMTs [15]. Some of these proteins might correspond to these inner proteins of the C2 microtubule. FAP65, 70, 75, 147, 171 and 239 were previously proposed to be somewhere at the C2 microtubule[37] and we were able to further narrow down the localizations of these proteins. For FAP123, 244 and 266, there was no previous information about their localization. FAP99 was previously assigned to the C1 microtubule based on the solubility but there was no direct evidence of localization. Tagging to FAP99 and structural analysis by cryo-ET in the future work would reveal this point.

Though not apparent like other proteins, we also assigned some CP candidates into other sub-structures. FAP216 did not resemble other known CP proteins’ MS profiles, being decreased in both *pf6* and *cpc1* strains and missing in *pf16* and *pf15* strains (Fig. S3D). This could mean that FAP216 is a scaffold protein reaching from the C1e to the C1f region (Fig. 5 and Table 2). CHLREDRAFT_170023 was also assigned to the C1b/f area since it is reduced in *cpc1* strain result though not significant by ANOVA test (Fig. 5 and Table 2).

In our MS profile, some of the known CP proteins also showed MS profiles which were not readily assigned to certain classes. A traditionally known C1 protein, PP1A, was detected only in WT sample (Fig. S3D) and therefore we were not able to assign it to certain sub-structures of the CP. Calmodulin also did not show significant decrease in either of the strain used in our study (Fig. S3D). As mentioned, Calmodulin is known to be shared with the RS. Thus, the decrease of Calmodulin is thought to be masked by the signal from RS. Similarly, FAP174 which is a traditionally known CP protein did not show significant decrease in either of strain by ANOVA test (Fig. S3D). Zhao *et al*., also failed to confidently assign this protein to certain location, but they found that FAP174 was immunoprecipitated by FAP246 which localizes to the C1b/f area [37]. In our MS profile, FAP174 indeed showed a slight decrease in *cpc1* strain which lacks the C1b/f region. Based on these results, FAP174 is thought to be located at the C1b/f area (Fig. 5 and Table 2) along with other compartments of axoneme. Phosphoglycerate mutase was also mapped to the C1b/f area since this protein is known to work with enolase which was described to localize at this sub-structure (Fig. 5 and Table 2). There were some remaining CP candidates which we were unable to assign to certain CP sub-structures (Fig. S3D and Table 2). Most of these proteins were, however, detected in small amounts and could be the result of false positives. Nonetheless, our comprehensive MS results using *Chlamydomonas* strains presented new CP protein candidates and information about the localizations for traditionally known and new CP proteins. These results will be a foundation for future studies focusing on obtaining the complete CP architecture utilizing protein tagging and cryo-ET.

From our MS profile, we were able to build a more complete model of the localizations of the CP proteins (Fig. 5). Our model of localizations also gave some insights into regulations of flagellar motility. FAP125 is a kinesin-like protein newly identified as a CP protein and localized to the C1c area based on our results. The presence of FAP125 at the C1c area is interesting since KLP-1, another known CP kinesin, is located at the C2c/d which is at the opposite side of the CP complex. KLP-1 was proposed to work as a conformational switch in CP[16], and thus, symmetrical binding of two CP-kinesins onto separate singlet microtubules might play a role in waveform switching or planar waveform in a coordinated way. Further functional analysis of FAP125 in future work will reveal this point. Like this example, our model of localizations of CP proteins can be used to understand how each CP protein is organized and working as a complex.

Our results also suggest the existence of interactions between CP proteins across CP sub-structures. DPY30, MOT17 and FAP81 were shown to localize at different CP sub-structures from our MS results (Fig. 5). These proteins, however, were shown to form a complex by immunoprecipitation result[37]. This suggests that larger super-complexes are formed by protein-protein interactions between neighboring sub-structures rather than several independent small sub-structures are attached to the CP singlets. Structural analysis of CP complex in a higher resolution would further reveal this point.

## Conclusion

PCD is a rare but prevalent congenital disease that derives from the impairment of motile cilia. An insufficient understanding of the protein composition of axonemal complexes directly affects the success and efficiency of clinical diagnosis of a wide variety of ciliopathies. By using comprehensive MS analysis of *Chlamydomonas* strains, we have identified novel proteins and localized them to specific sub-structures of the CP which allows for more informed interpretation of whole exome sequence data and cross-sectional analysis. Through this method, we circumvent traditional means of protein identification and localization and provide a more comprehensive insights into the entire making of the CP complex. Such proteomic approach by exploiting mutant strains would also be applicable for other uninvestigated areas of the axoneme. The novel proteins identified in this study also make for ideal candidates for further investigation of clinical researches for PCD.

## Acknowledgements

We thank Dr. Kaustuv Basu at the Facility for Electron Microscopy Research of McGill University for help in microscope operation and data collection. We are indebted to Mr. Lorne Taylor and Ms. Amy Ho Yee Wong from MUHC for help for mass spectrometry. This work was supported by grants from the Natural Sciences and Engineering Research Council of Canada (Discovery Grant 69462), Canada Institute of Health Research (Project Grant 388933) and the Canada Institute for Advanced Research Arzieli Global Scholars Program and McGill University to KHB. MI is supported by JSPS Overseas Research Fellowships.

## Author Contributions

MI and KHB conceived the project and designed the experiments. DD and MI performed culture of the cells, purification of microtubule fractions from flagella for MS analysis with the help of KP and RR. MI performed cryo-EM observation with the aid of KHB. DD and MI analyzed the results. All authors were involved in the manuscript writing process.

---

**Supplementary Table 1.**
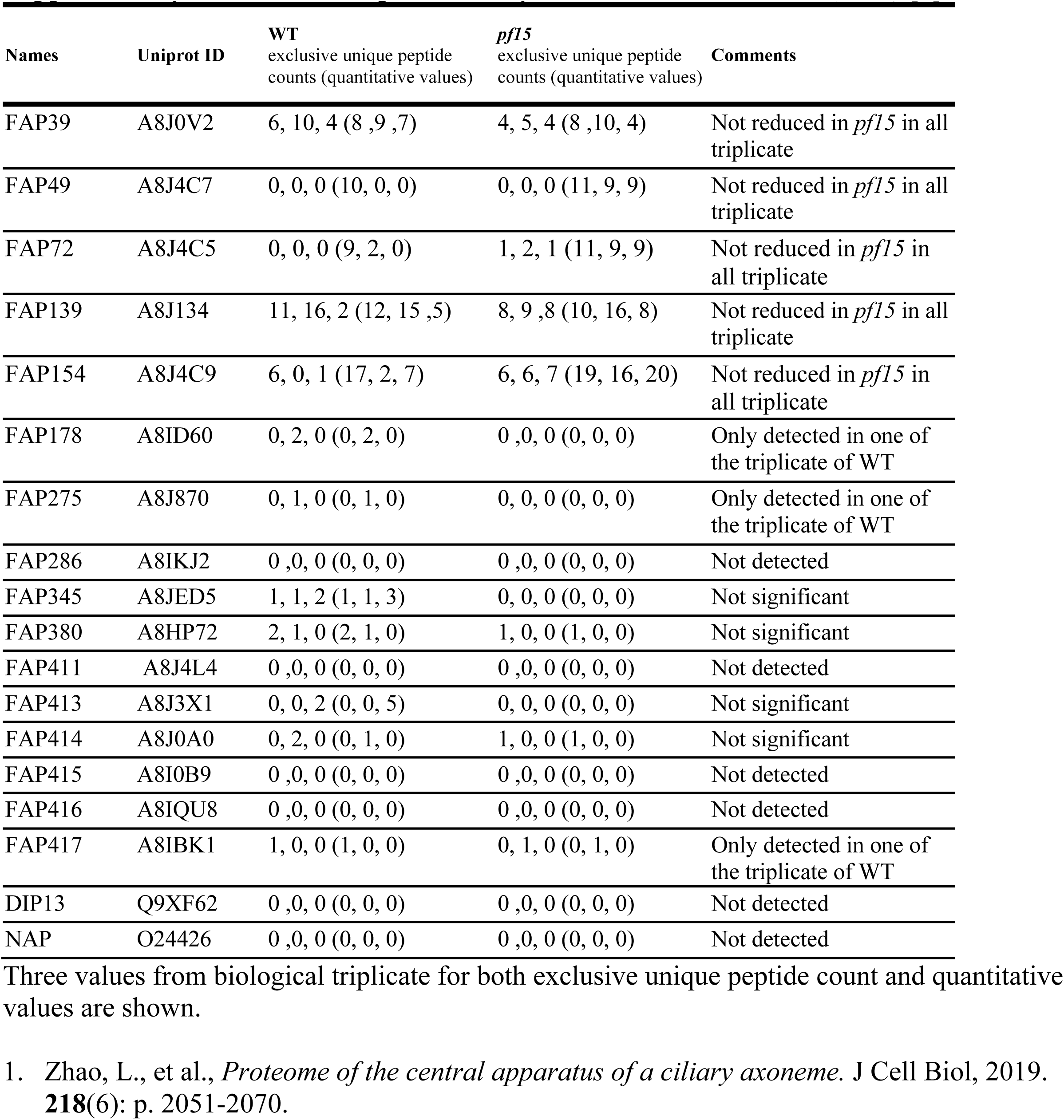
List of proteins only identified in Zhao *et al*., (2019) [1].

**Supplementary Fig. 1:**
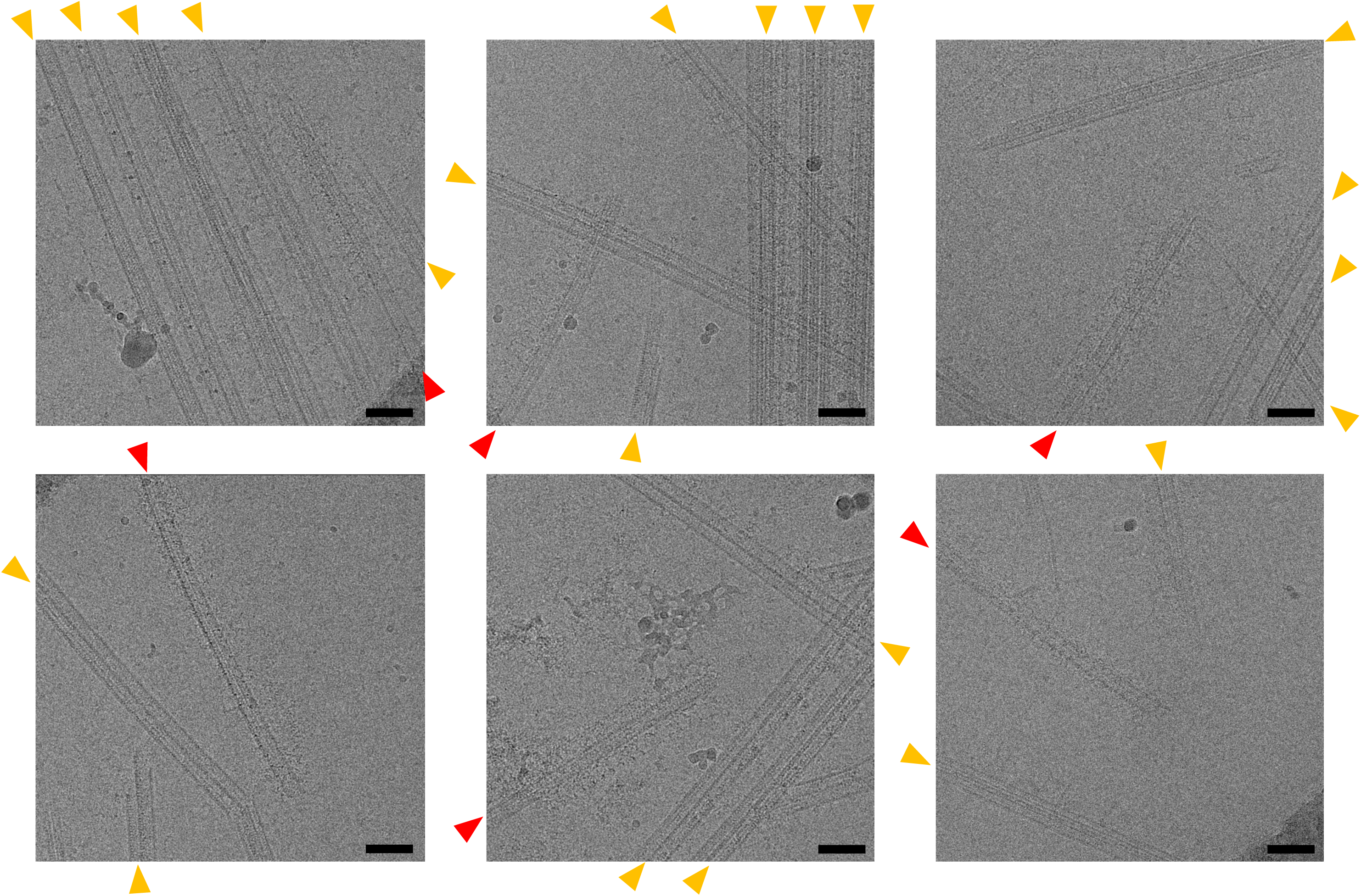
Additional cryo-EM images showing singlets from the CP. In our purified microtubule fractions, we occasionally observed singlet microtubules from the CP (red arrowheads) with characteristic appendages along with the DMTs (orange arrowheads). Scale bar, 100 nm.

**Supplementary Fig. 2:**
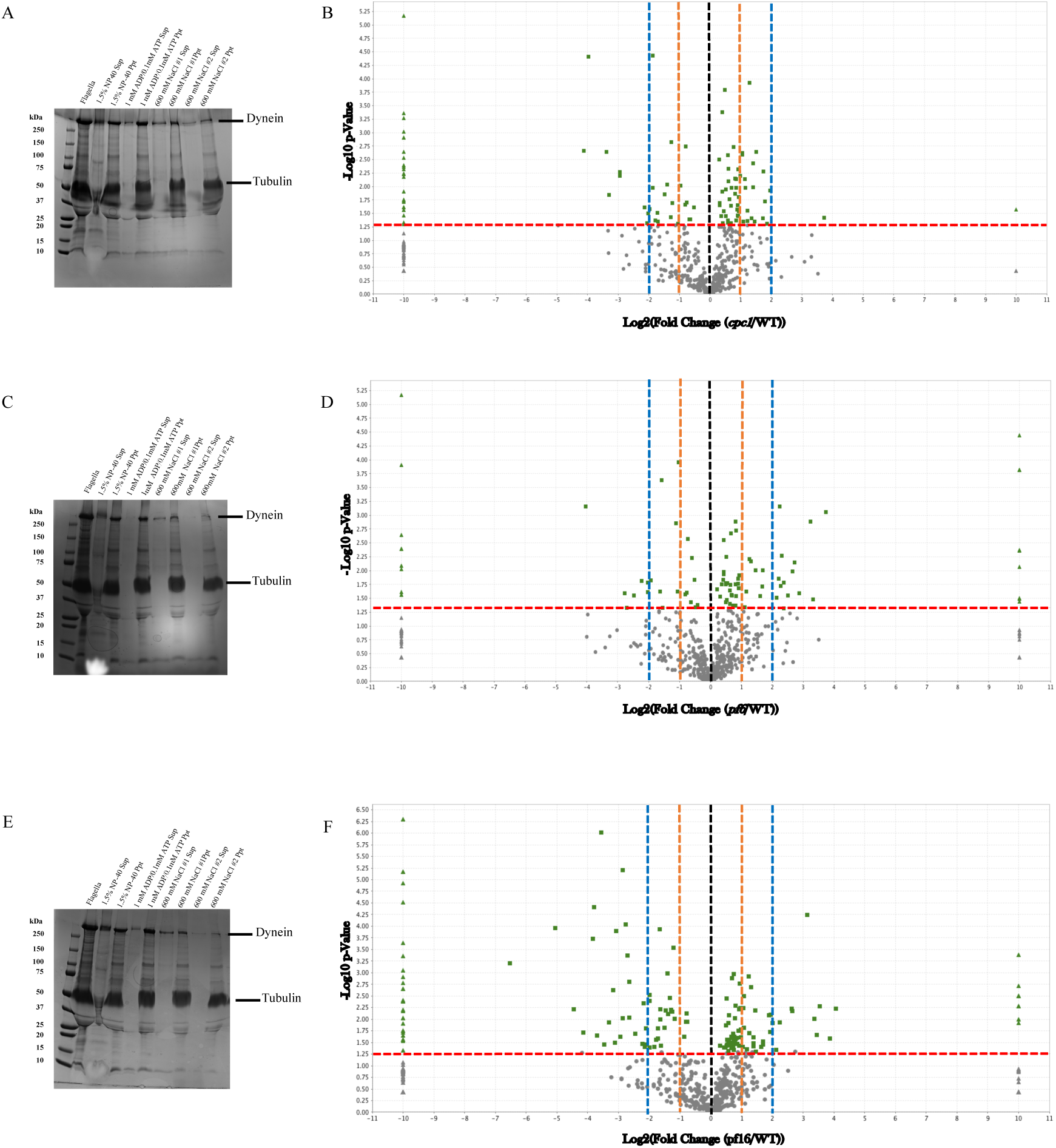
Purification of microtubule fractions from *cpc1, pf6* and *pf16* strains, and comparison of MS results with WT. (A and B) SDS-PAGE result of sequential purification of microtubule fraction from *cpc1* flagella and a volcano plot comparing *cpc1* result with WT result. (C and D) SDS-PAGE gel of purification from *pf6* axoneme and a volcano plot of *pf6* MS result compared with WT result. (E and F) *pf16* strain SDS-PAGE result and its MS result on a volcano plot compared with WT. Dashed red line indicates the significance threshold (*p* < 0.05) and proteins showed significant change are shown in green. Triangle dots represent completely missing proteins in either mutant results or WT result. Orange and blue dashed lines indicate two-and four-fold changes, respectively.

**Supplementary Fig. 3:**
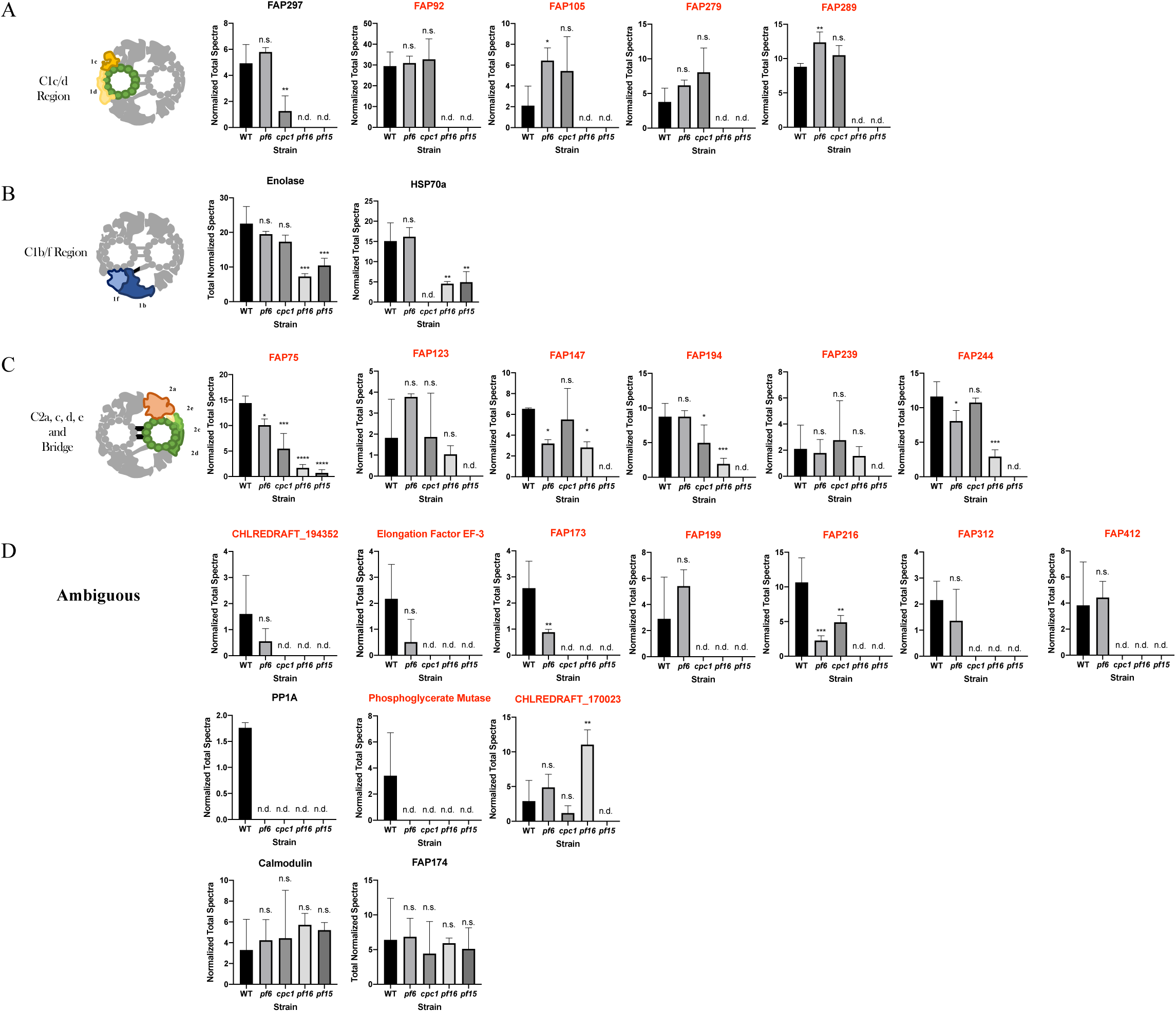
Additional MS profiles for other CP proteins. (A) MS profiles of proteins previously proposed to be localized at the C1d region and new CP proteins assigned to the C1d area. FAP297 was previously proposed to be a component of the C1d sub-structure, but its MS profile was different from other known C1d proteins and FAP297 is thought to be localized at a slightly different location. (B) MS profiles of CP proteins known to localize at the C1b/f area. HSP70 and enolase are known to be localized at the C1b/f, but MS profiles of these proteins were different from other known C1b/f proteins, possibly because these proteins are also shared with other axonemal complexes. (C) MS profiles of other new CP proteins mapped to the C2a, c, d, e and bridge area. (D) MS profiles of CP proteins which were not readily assigned to certain regions. Known CP protein Calmodulin, which is also shared with RS did not show significant change in either of strain tested here. Known CP protein FAP174 also showed similar trend with Calmodulin and thus thought to be shared with other axonemal complexes. Traditionally known CP proteins are shown in black and new CP proteins are shown in red through this figure.

